# To include or not to include: The impact of gene filtering on species tree estimation methods

**DOI:** 10.1101/149120

**Authors:** Erin K. Molloy, Tandy Warnow

## Abstract

Species tree estimation from loci sampled from multiple genomes is now common, but is challenged by the heterogeneity across the genome due to multiple processes, such as gene duplication and loss, horizontal gene transfer, and incomplete lineage sorting. Although methods for estimating species trees have been developed that address gene tree heterogeneity due to incomplete lineage sorting, many of these methods operate by combining estimated gene trees and are hence vulnerable to gene tree quality. There is also the added concern that missing data, which is frequently encountered in genome-scale datasets, will impact species tree estimation.

Our study addresses the impact of gene filtering on species trees inferred from multi-gene datasets. We address these questions using a large and heterogeneous collection of simulated datasets both with and without missing data. We compare several established coalescent-based methods (ASTRAL, ASTRID, MP-EST, and SVDquartets within PAUP*) as well as unpartitioned concatenation using maximum likelihood (RAxML).

Our study shows that gene tree error and missing data impact all methods (and some methods degrade more than others), but the degree of incomplete lineage sorting and gene tree estimation error impacts the absolute and relative performance of methods as well as their response to gene filtering strategies. We find that filtering genes based on the degree of missing data is either neutral or else reduces the accuracy of all five methods examined, and so is not recommended. Filtering genes based on gene tree estimation error shows somewhat different trends. Under low levels of incomplete lineage sorting, removing genes with high gene tree estimation error *can* improve the accuracy of summary methods, but only if not too many genes are removed. Otherwise, filtering genes tends to increase error, especially under high levels of incomplete lineage sorting. Hence, while filtering genes based on missing data is not recommended, there are conditions under which removing high error gene trees can improve species tree estimation. This study provides insights into prior studies and suggests approaches for analyzing phylogenomic datasets.

---

Species tree estimation is greatly enabled through the use of multiple loci, and increased access to genomic data over the last decade has opened up the possibility of greatly improving our understanding of how life has evolved on earth (Posada 2016). Furthermore, because species trees provide a context in which to understand evolution, including how genes evolve and how species adapt to their environments, improved species tree estimation can result in more accurate downstream analyses.

The estimation of species trees is complicated by biological processes, such as gene duplication and loss, horizontal gene transfer, and incomplete lineage sorting, resulting in different genomic regions, called “c-genes,” having different evolutionary histories (Ohno 1970; Syvanen 1985; Maddison 1997). Incomplete lineage sorting (ILS), which is modeled by the Multi-Species Coalescent (MSC) model (Pamilo and Nei 1988; Maddison 1997; Rannala and Yang 2003), is likely to occur with high frequency in the presence of rapid radiations, such as have occurred for the avian clades (Jarvis et al. 2014).

Several statistically consistent methods for inferring species trees under the MSC model have been developed and are increasing in use. Coalescent-based methods that combine estimated gene trees into a species tree, referred to as “summary methods”, include STAR (Liu et al. 2009), STEM (Kubatko et al. 2009), MP-EST (Liu et al. 2010), NJst (Liu and Yu 2011), iGLASS (Jewett and Rosenberg 2012), ASTRAL (Mirarab et al. 2014b; Mirarab and Warnow 2015), and a modification of NJst called ASTRID (Vachaspati and Warnow 2015). Statistical methods that co-estimate gene trees and species trees under the MSC model, such as *BEAST (Heled and Drummond 2010), are widely considered to be the most accurate methods, but they are computationally intensive on datasets with much more than 20 species and 50 genes (McCormack et al. 2009; Leavitt et al. 2016). BBCA (Zimmermann et al. 2014), which is a combination of *BEAST, random partitioning of genes into small disjoint bins, and summary methods, is an approach for improving the scalability of *BEAST to large numbers of genes; however, the BBCA technique does not improve the scalability of *BEAST to large numbers of species. Finally, site-based methods, such as SVDquartets (Chifman and Kubatko 2014, 2015) and SNAPP (Bryant et al. 2012), bypass gene tree estimation, inferring the species tree directly from the site patterns. These site-based methods have the potential to be more robust to low phylogenetic signal, and are increasingly popular.

The traditional approach to multi-locus species tree estimation is concatenation, in which the gene sequence alignments are combined into a single supermatrix, and then a phylogeny estimation method, e.g., RAxML (Stamatakis 2014), is used to construct the tree. Unpartitioned concatenation using maximum likelihood is commonly used for species tree inference but is not statistically consistent under the MSC model, and may even converge to the incorrect tree as the number of genes increases under some conditions with high levels of ILS (Roch and Steel 2015).

Because unpartitioned concatenation using maximum likelihood is statistically inconsistent under the MSC model, there is increasing interest in coalescent-based methods for species tree estimation, and especially in summary methods, because they are able to analyze large datasets. However, it is now well documented that high gene tree estimation error reduces the accuracy of species trees estimated using summary methods (Huang et al. 2010; Patel et al. 2013; Bayzid and Warnow 2013a; DeGiorgio and Degnan 2014; Mirarab et al. 2014a; Lanier and Knowles 2015; Mirarab and Warnow 2015; Xi et al. 2015; Meiklejohn et al. 2016). It is unknown whether summary methods remain statistically consistent when gene trees are not guaranteed to be accurate (Roch and Warnow 2015). Hence, there is substantial concern about the validity of estimating species trees using summary methods when gene trees have poor accuracy (Gatesy and Springer 2014; Springer and Gatesy 2016). Species tree error can also increase when genes are missing taxa, especially when datasets have limited numbers of genes (Hovmöller et al. 2013; Vachaspati and Warnow 2015; Xi et al. 2016), and it is unknown whether standard coalescent-based methods for species tree estimation remain statistically consistent under these conditions. Finally, little (if anything) is yet known about the response of SVDquartets to low phylogenetic signal and/or missing data (although see Chou et al. (2015b) for a study evaluating SVDquartets and some summary methods under changing numbers of sites per gene).

The impact of missing data on concatenation-based methods is well-studied on both biological and simulated datasets (Wiens 2006; Lemmon et al. 2009; Wiens and Morrill 2011; Roure et al. 2013; Simmons 2014). Some studies (Wiens and Morrill 2011; Lemmon et al. 2009; Simmons 2014) have examined the impact of missing data on simulated multi-gene datasets with branch lengths (but not tree topology) varying across genes. Although Wiens and Morrill (2011) find missing data to be unproblematic provided that each taxon has sufficiently many characters in the data matrix, other studies observe systematic biases due to missing data under certain model conditions, e.g., when the distribution of missing data is non-random (Simmons 2014) or when the evolutionary rates vary greatly across genes (Lemmon et al. 2009). The mixed results of these studies may have led to some phylogenomic analyses using concatenation to remove taxa (McKenna et al. 2015) or genes (Pyron et al. 2014) with high degrees of missing data.

Perhaps because prior simulation studies have established that gene tree estimation error and missing data can reduce species tree accuracy, filtering genes prior to species tree estimation is very common in both concatenation-based and coalescent-based analyses of biological datasets (Jiang et al. 2014; Chen et al. 2015; Hosner et al. 2016; Simmons et al. 2016; Streicher et al. 2016; Blom et al. 2017). However, the ability of empirical studies to assess the impact of gene filtering on species tree accuracy is limited (i.e., the true species tree is rarely known with certainty); instead, empirical studies typically approach this issue by examining the recovery of well-established clades and their bootstrap support, or the stability of the analysis (i.e., the similarity between the tree constructed on the full set of genes compared to the tree constructed on the subsets of genes). As an example, Jiang et al. (2014) use similarity to the concatenation analysis on the full dataset to evaluate the impact of filtering based on missing data. Although their study was limited to two biological datasets (yeasts and plants) and only evaluated the impact of filtering on concatenation analyses using maximum likelihood, their study does suggest that concatenation analyses decrease in accuracy with filtering based on degrees of missing data. They argue that further work is needed to understand the impact of filtering on other methods (and especially on current species tree methods that are designed to address gene tree heterogeneity). The questions of whether gene filtering should be performed, and if so how and under what circumstances, are among the current important methodological questions in phylogenomic estimation.

To the best of our knowledge, only one prior study (Huang and Knowles 2016) has examined the impact of gene filtering on datasets simulated under the MSC model. They specifically examine the question of gene filtering based upon the degree of missing data, using a simulation designed to produce datasets similar those generated by RADtag protocols (Baird et al. 2008). For these simulated datasets, Huang and Knowles (2016) find that filtering genes with missing data has the unfortunate consequence of removing genes with higher rates of evolution. They noted that these deleted genes are the ones that provide resolution at difficult nodes, so that the consequence of deleting these genes is *an increase in species tree estimation error*.

Thus, Huang and Knowles (2016) provide evidence that screening genes on the basis of missing data can reduce accuracy of estimated species trees. However, their simulation study was specifically designed to mimic properties of RADseq datasets, and so is not immediately generalizable to other types of sequencing protocols where rates of evolution are not necessarily correlated with the probability of missing data (Fig. S1). In addition, their study computed species trees using the shallowest divergence method (Takahata 1989), a method that is similar in design to STAR in that it estimates distances between species for each gene and uses these estimates to construct the species tree. Therefore, their observations may not apply to other methods not based on estimating pairwise distances. Finally, their study was limited to datasets with only 8 species, and may not be generalizable to larger datasets.

We report on a simulation study in which we examine the impact of gene filtering based on either missing data or on gene tree estimation error (a question that seems to not have been addressed in any prior simulation study). Our simulation protocol differs from Huang and Knowles (2016) in that the missing data is biased towards a subset of genes but the probability of a gene being deleted from a species is independent of its rate of evolution. Our study addresses the following questions:

1. How does gene tree estimation error and degree of ILS impact the absolute and relative accuracy of different methods?
2. How does gene filtering based on gene tree estimation error impact species tree accuracy?
3. How does gene filtering based on missing data impact species tree accuracy?

Our study shows that different species tree estimation methods respond differently to different model conditions, and that gene tree estimation error and ILS level are critically important to the impact of gene filtering. The prediction of gene tree estimation error is typically assessed using some way of measuring statistical support (e.g., average bootstrap support on the branches), which seem to be reasonable proxies for gene tree accuracy when the gene sequence evolution is a reasonable match to the estimation model (Anisimova et al. 2011). However, much less is understood about how to predict the ILS level from empirical data. Therefore, we also address the following question:

1. How well does heterogeneity between the species tree and estimated gene trees, a metric commonly reported in biological studies, reflect the level of heterogeneity due to ILS?

We address the first three questions using four methods designed to address gene tree heterogeneity due to ILS: three summary methods (ASTRAL, ASTRID, and MP-EST) and a site-based method (SVDquartets within PAUP* (Swofford 2016)), which has been identified as a promising new approach that may provide better accuracy than summary methods under conditions where gene trees have low accuracy. We also include concatenation under maximum likelihood, where gene alignments are concatenated into a single “supermatrix” and then unpartitioned RAxML is run on the concatenated matrix. Although this approach is not statistically consistent, it has been shown to outperform coalescent-based methods on some simulated datasets (Patel et al. 2013; Bayzid and Warnow 2013b; Mirarab and Warnow 2015; Chou et al. 2015a), and so is an important technique to examine.

We find that unpartitioned concatenation using RAxML outperforms the other methods under many conditions. When the level of ILS is fairly high, summary methods (ASTRAL, ASTRID, and MP-EST) can be more accurate than concatenation provided that the level of gene tree estimation error is sufficiently low. When gene tree estimation error is high, SVDquartets is more accurate than the summary methods but otherwise is often among the least accurate methods.

Responses to gene filtering (where genes are removed from the dataset because they have too much missing data or too high gene tree estimation error) also depend on model condition and method. Filtering genes due to gene tree estimation error can improve accuracy for summary methods, but only when ILS is sufficiently low – otherwise it reduces accuracy. SVDquartets and concatenation become less accurate with gene filtering based on gene tree estimation error; the explanation is that they do not utilize gene trees and hence do not benefit from the removal of gene sequences corresponding to trees with high estimation error. Filtering genes due to missing data generally reduces accuracy for all methods, and so is not recommended.

As the levels of ILS and gene tree estimation error impact the relative method performance and the impact of filtering, the ability to estimate these two factors can inform the selection of phylogenomic species tree estimation methods for empirical datasets. However, our study also shows that a standard approach for estimating the ILS level that compares estimated gene trees to a reference or estimated species tree can greatly inflate the ILS level whenever gene tree estimation error is not low. Hence, the development of techniques to estimate ILS levels in empirical data is an important area of research.

## MATERIALS AND METHODS

### Overview

Our study evaluates three summary methods (ASTRAL v4.10.5, ASTRID v1.1, and MP-EST v1.5), one site-based coalescent method (SVDquartets within PAUP* v4a152), and concatenation using RAxML v8.2.8. We compare these methods on a collection of 26-taxon simulated datasets derived from Mirarab and Warnow (2015). These datasets provide a range of model conditions from relatively easy (e.g., low levels of gene tree estimation error and ILS) to very challenging (e.g., high levels of gene tree estimation error and ILS), both with and without missing data. We explore the impact of gene filtering on species tree estimation, where genes are filtered on the basis of missing data or gene tree estimation error. Distances between trees are measured using the normalized Robinson-Foulds (Robinson and Foulds 1981) tree distance. Our study estimated 480,000 gene trees and 14,400 species trees, requiring ~3,000 CPU hours or over 4 months of CPU time.

### Simulated Datasets

The datasets for this study were originally simulated by Mirarab and Warnow (2015). Because MP-EST is computationally intensive on datasets with even 50 species (Bayzid et al. 2014; Mirarab and Warnow 2015), we restricted the 200-taxon datasets to 26 taxa (the outgroup taxon and 25 randomly selected taxa) and used only 20 (out of the original 50) replicates in our simulation study.

#### Incomplete Lineage Sorting

Mirarab and Warnow (2015) simulated datasets using SimPhy (Mallo et al. 2016) with three species tree heights (10M, 2M, and 500K generations) for both deep (rate: 10^−7^) and recent (rate: 10^−6^) speciation. As the population size was fixed across all model conditions, shorter tree heights correspond to higher levels of ILS. The ILS level for each model condition is measured by the average distance (AD) between the true species tree and true gene trees across all 1000 genes. The mean AD (± standard deviation) across replicates corresponding to *low/moderate* ILS (species tree height: 10M), *high* ILS (species tree height: 2M), and *very high* ILS (species tree height: 500K) is 12 ± 2%, 41 ± 6%, and 75 ± 1%, respectively.

#### Gene Tree Estimation Error

Mirarab and Warnow (2015) simulated gene sequence data under a GTR+Γ model of evolution from true gene trees with branch lengths deviating from a strict molecular clock. Sequence lengths were drawn from a distribution varying from 300 to 1500 sites. We truncated sequences to the first 100 base pairs to mimic conditions where sequences are shortened to avoid recombination (Hobolth et al. 2011) or sequences are drawn from highly conserved regions of the genome having very few variable sites (Hosner et al. 2016).

This process produced datasets with a range of GTEE consistent with the bootstrap support values seen in biological datasets. For example, the avian phylogeny computed in Jarvis et al. (2014) was based on over 14,000 markers, including exons, introns, and UCEs. The average bootstrap support of the gene trees on these markers was generally low: exons had average boostrap support of 26%, UCEs had average bootstrap support of 40%, and introns had average bootstrap support of 47% (Table 1). Table 1 presents similar statistics for three other biological datasets. Thus, phylogenomic analyses, especially when performed on genome-scale data, are likely to be based on genes that have very low average bootstrap support. Under the reasonable assumption that the low bootstrap edges are less likely to be accurate, this suggests that the typical gene tree in a large-scale phylogenomic analysis will have high error rates.

**Table 1:** Empirical statistics from biological datasets are shown. We report mean bootstrap support (BS), defined as the bootstrap support of the bootstrap greedy consensus (GC) gene tree averaged across all branches; the normalized Robinson-Foulds (RF) distance between the bootstrap GC gene trees and the concatenation-based analysis using maximum likelihood (CA-ML) species tree; and the normalized False Positive (FP), False Negative (FN), and RF distances between the bootstrap Strict Majority Consensus (SMC) gene trees and the CA-ML species tree. Means and standard deviations are across all loci.

| Study | Dataset Type | Number of Taxa | Number of Loci | Mean BS of GC | Distance between GC and CA-ML | Distance between SMC and CA-ML |  |  |
| --- | --- | --- | --- | --- | --- | --- | --- | --- |
|  |  |  |  |  |  | FP | FN | RF |
| Blom et al., 2017 | Exon | 29 | 1361 | $0.28 \pm 0.11$ | $0.79 \pm 0.12$ | $0.45 \pm 0.26$ | $0.87 \pm 0.09$ | $0.80 \pm 0.13$ |
| Hosner et al., 2016 | UCE | 91 | 4817 | $0.28 \pm 0.12$ | $0.74 \pm 0.20$ | $0.37 \pm 0.25$ | $0.83 \pm 0.17$ | $0.76 \pm 0.19$ |
| Jarvis et al., 2014 | Exon | 48 | 8251 | $0.26 \pm 0.07$ | $0.78 \pm 0.09$ | $0.22 \pm 0.25$ | $0.85 \pm 0.08$ | $0.75 \pm 0.12$ |
| Jarvis et al., 2014 | Intron | 48 | 2516 | $0.47 \pm 0.12$ | $0.62 \pm 0.10$ | $0.20 \pm 0.13$ | $0.68 \pm 0.09$ | $0.55 \pm 0.09$ |
| Jarvis et al., 2014 | UCE | 48 | 3679 | $0.40 \pm 0.05$ | $0.65 \pm 0.05$ | $0.11 \pm 0.11$ | $0.71 \pm 0.03$ | $0.57 \pm 0.04$ |
| Streicher et al., 2016 | UCE | 29 | 4784 | $0.39 \pm 0.09$ | $0.71 \pm 0.13$ | $0.36 \pm 0.26$ | $0.78 \pm 0.12$ | $0.69 \pm 0.16$ |

We estimated ML gene trees using RAxML v8.2.8 with a single tree search under the GTRGAMMA model of evolution. We use maximum likelihood gene trees rather than bootstrap gene trees, as this tends to result in improved accuracy of summary methods under many conditions (Mirarab et al. 2016).

Replicates (out of the original 20) were then partitioned based on their mean gene tree estimation error (GTEE), defined as the distance between the true and estimated gene trees averaged over across all genes. Specifically, replicates in the full length sequence datasets (300-1500 bp) were partitioned by *low/moderate* GTEE (i.e., mean GTEE within 0-20%) and *moderate/high* GTEE (i.e., mean GTEE within 20-50%). The mean GTEE averaged across replicates (± standard deviation) is 16±2% and 35±8% for *low/moderate* and *moderate/high* GTEE, respectively (Tables S2-S3). Replicates in the truncated sequence datasets (100 bp) were partitioned by *very high* GTEE (i.e., mean GTEE within 50-80%) and *extremely high* GTEE (i.e., mean GTEE within 80-100%). The mean GTEE averaged across replicates (± standard deviation) is 69 ± 8% and 86 ± 5% for *very high* and *extremely high* GTEE, respectively (Tables S4-S5).

### Gene Filtering Experiments

Filtering out low quality gene trees comes at the cost of fewer genes, and increasing the number of gene trees has been shown to make methods more robust to GTEE (Xi et al. 2016) as well as to missing data (Xi et al. 2015). We evaluate the impact of gene filtering by removing 0%, 25%, 50%, 75%, 90%, and 95% of the genes, thus producing datasets that vary in the number of genes retained for species tree inference.

#### Gene Tree Estimation Error (GTEE)

Gene trees are sorted based on GTEE and 0%,25%, 50%, 75%, 90%, and 95% of the genes with the highest GTEE are removed prior to species tree estimation.

#### Missing Data

We deleted genes from species using a protocol designed to produce datasets with a similar total amount and distribution of missing data as Hosner et al. (2016) dataset (supplement). To missing data biased towards a random subset of genes, 250 genes were selected at random. The exact number of missing taxa was determined by drawing a random value between 13 and 19, and sequences were deleted at random from the gene alignment, producing a gene with between 50% and 73% missing data. This protocol was repeated to produce genes with varying degrees of missing data. Specifically, 250 genes are missing between 7-12 sequences (i.e., 27-46%), 250 are missing between 3-6 sequences (i.e., 12-23%), 100 genes are missing 2 sequences (i.e., 8%), 50 genes are missing 1 sequence (i.e., 4%), and 50 genes are missing no sequences.

The total amount of missing data is approximately 30% for all datasets, but this protocol produces genes where some are missing a large number of species and other genes are missing a small number of species; this produces a pattern that is similar to the Hosner et al. (2016) dataset. In addition, this protocol creates a condition where removing genes missing at least 50%, 25%, 10%, or 5% of the species results in datasets with the same number of genes as filtering based on GTEE, making these two experiments directly comparable.

### Species Tree Estimation

Five methods for species tree estimation methods are evaluated: three gene tree summary methods (ASTRAL v4.10.5, ASTRID v1.1, and MP-EST v1.5), the site-based method SVDquartets using PAUP* v4a150, and concatenation under maximum likelihood (RAxML v8.2.8) under the GTRGAMMA model on the unpartitioned concatenated alignment. The summary methods are all run on the best maximum likelihood gene trees. ASTRAL and ASTRID are run in default mode on unrooted gene trees. Since MP-EST requires rooted gene trees, estimated gene trees are rooted at the outgroup when available, and otherwise rooted at the midpoint of the longest leaf-to-leaf path using Dendropy v4.1.0 (Sukumaran and Holder 2010). For MP-EST, the best pseudo-likelihood scoring species tree is taken from ten independent runs of MP-EST. Branch support in species trees estimated using ASTRAL is computed using local branch support, which is based on the quartet frequency estimated from the input set of gene trees (Sayyari and Mirarab 2016).

## RESULTS

### Species Tree Method Performance

Our first experiment evaluates methods on the full set of 1000 genes (i.e., without filtering) and shows that species tree error rates increase for all methods as ILS and/or GTEE levels increase (Fig. 1).

**Figure 1:**
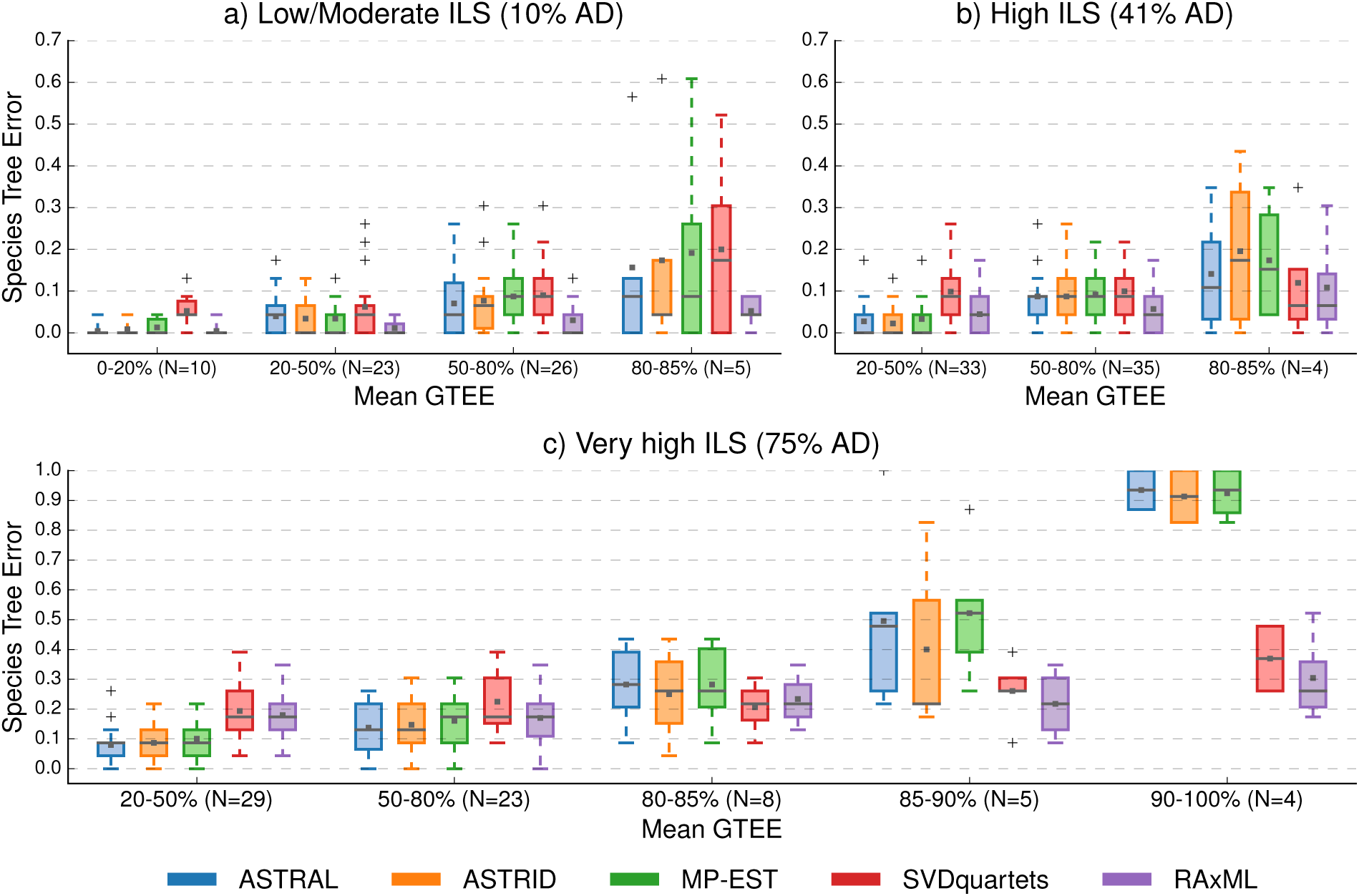
The impact of mean gene tree estimation error (GTEE) level and ILS level on species tree error is shown for five methods: ASTRAL (blue), ASTRID (orange), MP-EST (green), SVDquartets (red), and RAxML (purple). Species tree error is the normalized RF distance between the true and estimated species trees. Mean GTEE is the normalized RF distance between the true and estimated gene trees averaged across all genes. Subplots a, b, and c show three levels of increasing ILS, where average distance (AD) is the normalized RF distance between the true species and true gene trees averaged across all genes. The number of replicates (N) for each model condition is specified on the x-axis. Means and medians are denoted by the gray dot and bar, respectively. Box plots are defined by quartiles, e.g., boxes extend from the first to the third quartiles. Increases in ILS or GTEE result in increased species tree error rates for all methods, and relative performance depends on both ILS and GTEE. Under low to moderate ILS, RAxML tends to have better accuracy than the other methods, even for the highest GTEE condition. Under higher levels of ILS, summary methods are typically more accurate than RAxML and SVDquartets except for high GTEE conditions.

Under the low/moderate ILS condition, concatenation using RAxML is the most accurate method (Fig. 1(a)) for all levels of GTEE. When mean GTEE is less than 50%, all five methods have good accuracy with mean species tree error below 7%. Species tree error increases with GTEE, and the differences between methods are noteworthy. When mean GTEE is extremely high (80-85%), the mean species tree error rate for concatenation is 5%, the mean error rates for summary methods range from 16% (ASTRAL) to 19% (MP-EST), and the mean error rate for SVDquartets is 20%.

Under higher levels of ILS, the relative performance between methods changes dramatically with GTEE. Specifically, the three summary methods outperform SVDquartets and concatenation when GTEE is moderate (i.e., mean GTEE <50%), but SVDquartets and concatenation outperform the summary methods when GTEE is extremely high (i.e.,mean GTEE ≥80%) (Fig. 1(b-c)). For the highest level of ILS (75% AD) and the highest level of mean GTEE (90-100%), the differences in accuracy between the summary methods and site-based methods are very large: the mean species tree error rates for summary methods are all greater than 90%, but the error rates for concatenation and SVDquartets are 30% and 37%, respectively (Fig. 1(c)).

We also evaluate the impact of missing data (see protocol described above) on method accuracy (Fig. 2). Overall, the reduction in accuracy tends to be fairly low (typically below 5%) under most model conditions, although methods differ somewhat in their response to missing data. ASTRAL and ASTRID are quite robust with mean species tree estimation error never increasing by more than 6% and most increases in error are smaller. Under very high level of ILS, MP-EST increases by 10% under 80-85% mean GTEE, and both concatenation and SVDquartets both increase by 10% under 90-100% mean GTEE (Fig. 2(c)). These trends suggest that missing data, although generally detrimental, do not result in a huge increase in error, and that some methods become less robust to missing data when both ILS and GTEE levels are high.

**Figure 2:**
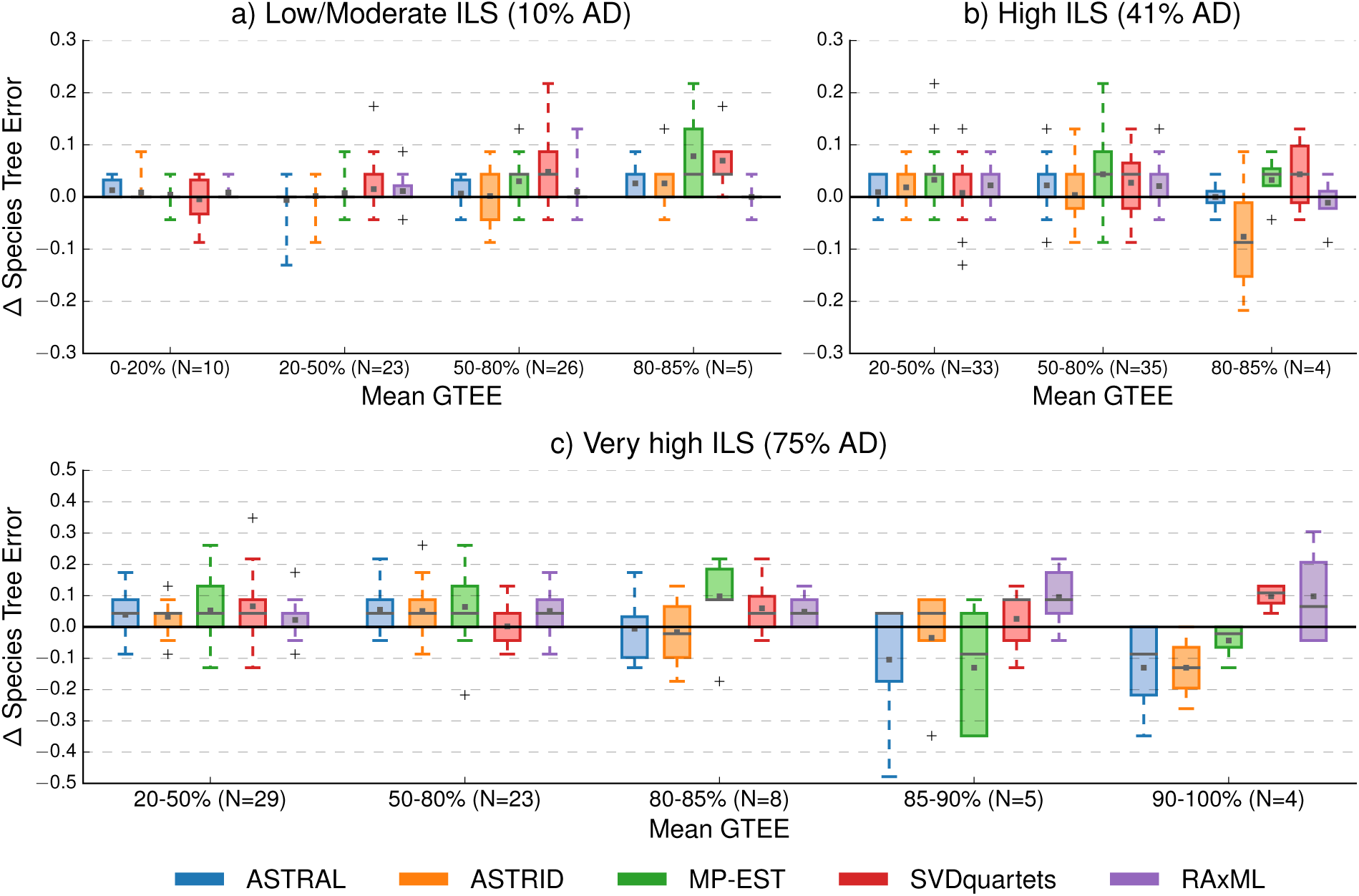
Differences in species tree error between datasets with no missing data and datasets with 30% missing data are shown for five methods: ASTRAL (blue), ASTRID (orange), MP-EST (green), SVDquartets (red), and RAxML (purple). Positive values indicate increases in error, whereas negative values indicate reductions in error. Subplots a, b, and c show three levels of increasing ILS. The number of replicates (N) for each model condition is specified on the x-axis. Means and medians are denoted by the gray dot and bar, respectively. Box plots are defined by quartiles, e.g., boxes extend from the first to the third quartiles.

There are even a few conditions, characterized by very high ILS and very high GTEE, where the accuracy of some summary methods *improves* on datasets with incomplete genes. The most likely explanation is that gene trees for those cases have extremely short branches (because species trees with very high ILS have very short branches), and the random deletion of taxa from gene trees may increase the branch lengths and make the gene trees easier to estimate. Thus, summary methods, which combine estimated gene trees, have the potential to return more accurate species trees.

Overall, trends in relative method accuracy are the same for datasets with and without missing data (Fig. S4).

### Filtering based on Gene Tree Estimation Error

The impact of filtering genes based on GTEE depends on the ILS level, with summary methods improving under filtering when the level of ILS is sufficiently low (12% AD), provided that the number of genes does not decrease too substantially (Fig. 3a-b). For example, the removal of ~75% of the genes based on GTEE results in ~2-3% improvement in species tree accuracy for the three summary methods under the lowest level of ILS. Under the higher ILS levels, filtering based on GTEE is at best neutral, but generally increases species tree error (Fig. 3(c)-(f)). Although RAxML and SVDquartets do not rely on estimated gene trees, they also decrease in accuracy when gene sequences corresponding to high error gene trees are removed (Fig. S5).

**Figure 3:**
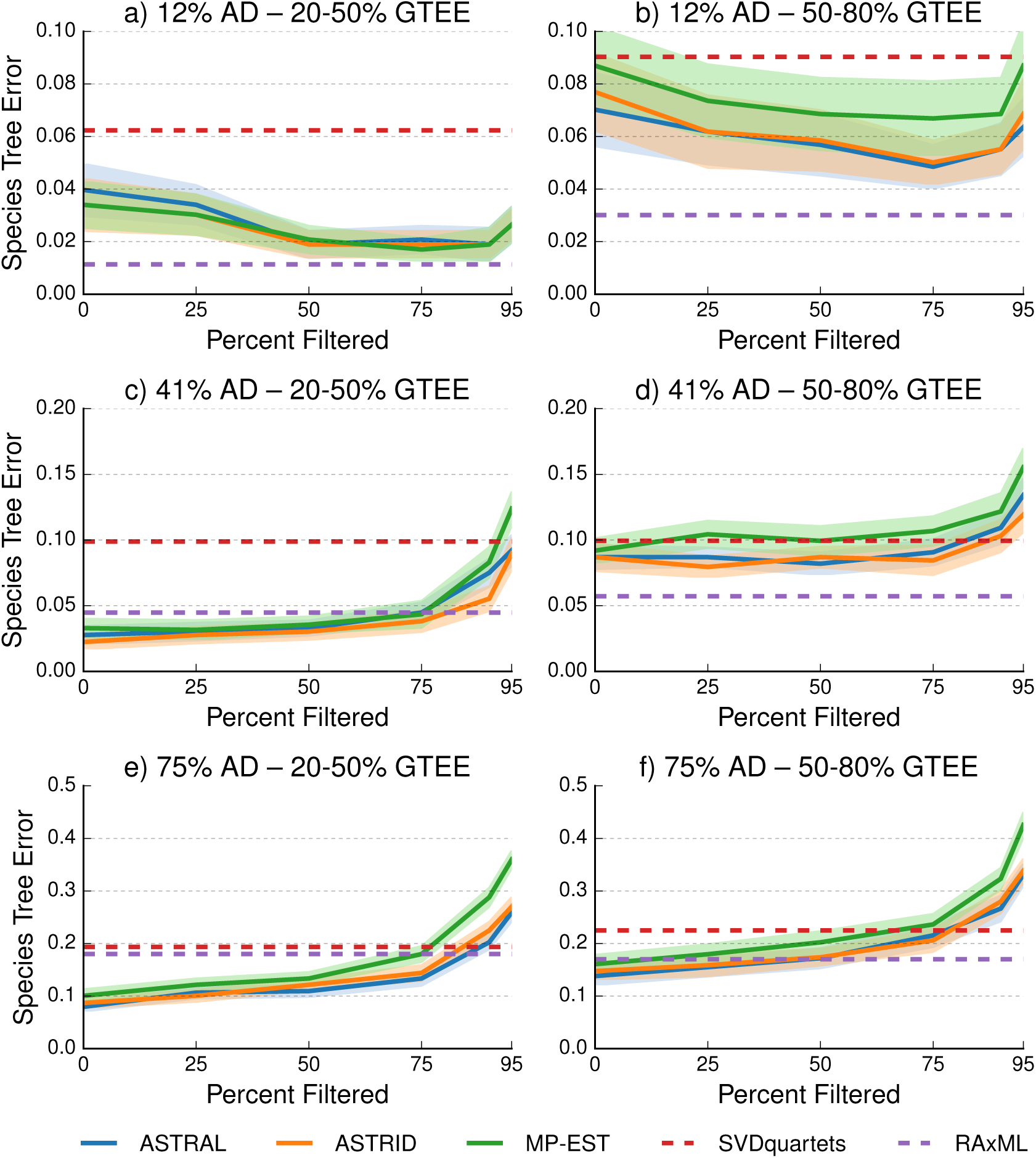
The impact of filtering genes by gene tree estimation error (GTEE) on species tree error is shown for three gene tree summary methods: ASTRAL (solid blue), ASTRID (solid orange), and MP-EST (solid green). Genes were filtered by removing 25%, 50%, 75%, 90%, and 95% of genes with the highest GTEE. Species tree error rates for SVDquartets (dashed red) and RAxML (dashed purpose) are shown on the unpartitioned concatenated alignment with no filtering. Lines indicate the mean across all replicates, and filled regions indicate the standard error. Rows show three levels of increasing ILS, where average distance (AD) is defined as the normalized RF distance between the true species tree and true gene trees averaged across all genes. Columns show two levels of GTEE. When ILS is sufficiently low, gene filtering (up to 75%) increases the accuracy of gene tree summary methods (a-b). When ILS is high to very high, gene filtering has no impact on the accuracy of gene tree summary methods or else reduces summary method accuracy (d-f).

Although summary methods can become more accurate with gene filtering based on GTEE for the lowest ILS condition examined (12% AD), unpartitioned concatenation using RAxML remains the most accurate method for this ILS level. Hence, although filtering genes based on GTEE can benefit summary methods, filtering does not make them as accurate as concatenation under these conditions.

Gene filtering based on GTEE has minimal impact on ASTRAL’s local branch support (Fig. S6-S7); however, it can decrease the mean accuracy of the true branches recovered by ASTRAL, increasing the number of true branches with low support (< 75%) by ~5%.

As expected, gene filtering based on GTEE reduces the average GTEE of the remaining genes as compared to the original set of genes (Fig. S8). Reductions in mean GTEE that can be very large. For example, when GTEE is moderate/high, the mean GTEE of the unfiltered datasets is 35-40%, while the mean GTEE is ~20% after the removal of 75% of the genes (Fig. S8 (a,c,e)).

### Filtering based on Missing Data

Filtering genes based on missing data is at best neutral, but frequently reduces the accuracy of estimated species trees; this trend holds regardless of the level of ILS or GTEE and for all the methods examined (Fig. 4). When the ILS and GTEE levels are sufficiently low, the impact of gene filtering tends to be modest, but the impact can be much larger under other conditions. For example, methods decrease in accuracy by ~5% as the number of genes is reduced from 1000 to 50 when the level of ILS is low/moderate (12% AD) and the level of GTEE is moderate/high (20-50%). When ILS and GTEE levels are both very high, accuracy decreases by at most 3% as the number of genes is reduced from 1000 to 500 (corresponding to removing genes with ≥50% missing data), and accuracy is decreased by ~25% as the number of genes decreases from 500 to 50 (corresponding to removing all remaining genes with missing data).

**Figure 4:**
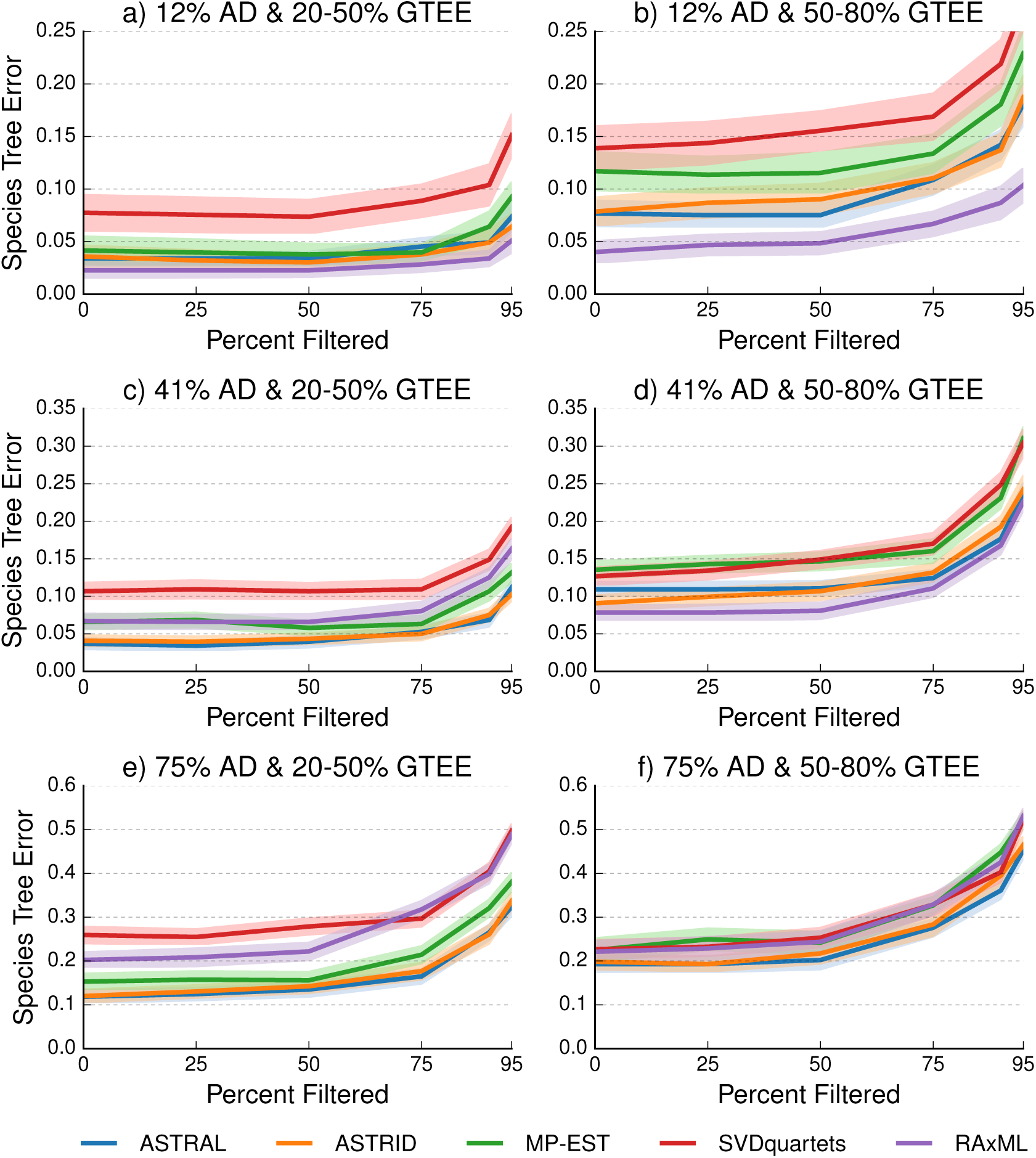
The impact of filtering genes by missing data on species tree error is shown for five methods: ASTRAL (blue), ASTRID (orange), MP-EST (green), SVDquartets (red), and RAxML (purple). Genes were filtered by ranking the 1000 genes based on the percentage of missing data and removing the 25%, 50%, 75%, 90%, and 95% of the genes according to this ordering; this resulted in removing genes with at least 50%, 25%, 10%, 5%, and 1% of missing taxa. Lines indicate the mean across all replicates, and filled regions indicate the standard error. Rows show three levels of increasing ILS, where average distance (AD) is defined as the normalized RF distance between the true species tree and true gene trees averaged across all genes. Columns show two levels of gene tree estimation error (GTEE). Gene filtering based on missing data is at best neutral, but often reduces the accuracy of species tree estimation methods.

Filtering genes based on missing data can decrease the mean branch support for true branches recovered by ASTRAL, resulting in a higher frequency of true branches with support less than 75% (Fig. S10-S11), and has a modest (but sometimes beneficial) impact on the mean GTEE of the retained set of genes (Fig. S12).

### Estimating ILS Levels

As the level of ILS levels impacts gene tree heterogeneity as well as method selection and the impact of gene filtering, we consider the practice of reporting the average RF distance between estimated gene trees and a reference (or estimated) species tree as an indication of the gene tree heterogeneity in a biological dataset (see, for example, the discussion in Jarvis et al. (2014)). We examine this metric using the datasets simulated for our study, as well as simulated datasets representative of those used in other studies (Fig. 5 and Supplement). The four subfigures correspond to different ranges of mean GTEE for estimated gene trees. Within each subfigure we show a scatterplot for the AD on the y-axis versus total discord (i.e., the normalized RF distance between estimated gene trees and the true species tree) on the x-axis. For the lowest ILS level, there is a nearly perfect relationship, so that total discord is an excellent predictor of the AD value (Fig. 5(a)). For high levels of GTEE, the difference between the total discord and the AD value increases, making the total discord a very poor predictor of the AD value (Fig. 5(b)-(d)). This experiment shows that except when gene tree estimation error is very low, much of the total discord is due to gene tree estimation error, making the total discord a poor estimate of the discord between true gene trees and the true species trees, which in these simulations is due entirely to ILS.

**Figure 5:**
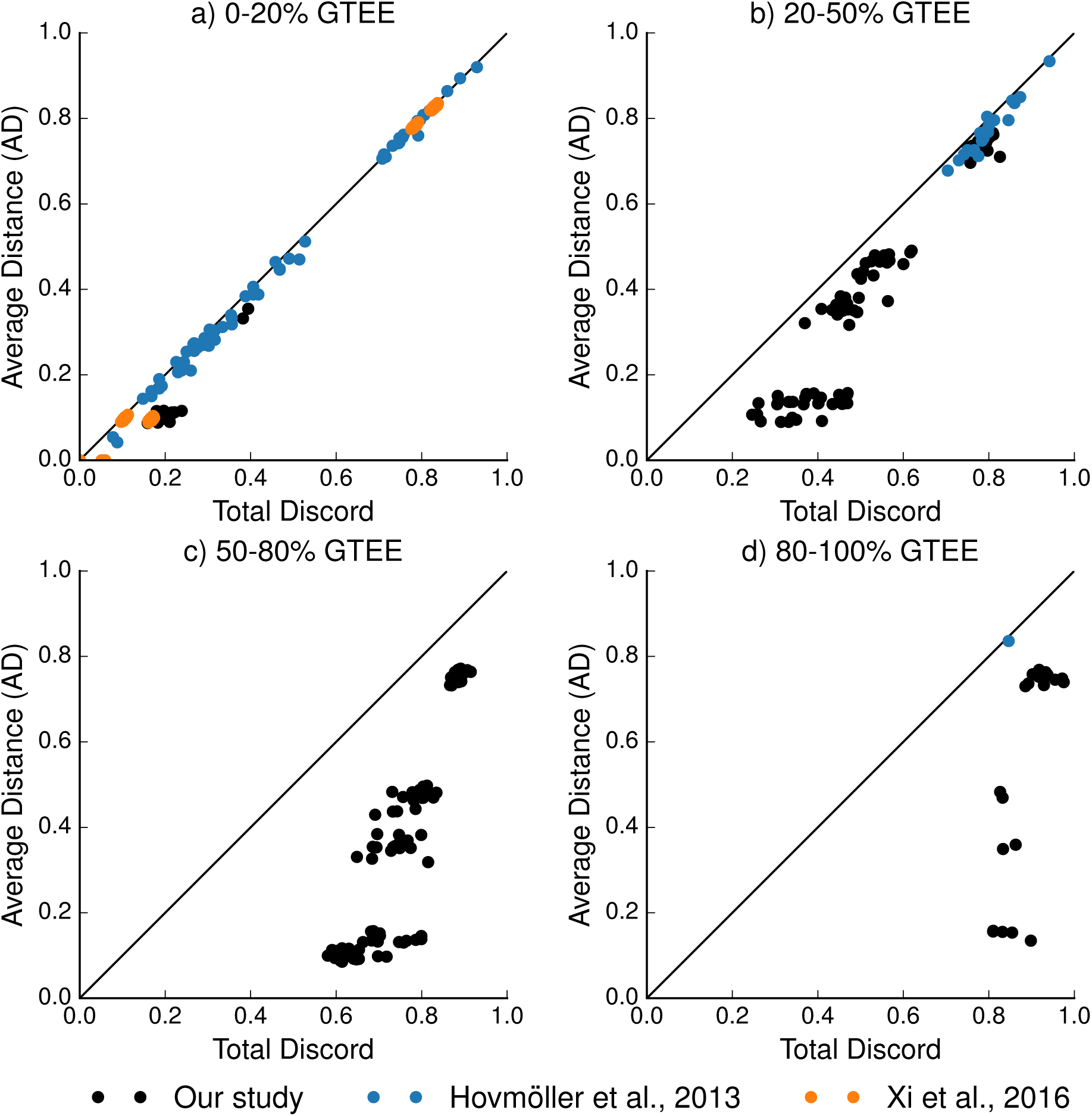
The level of incomplete lineage sorting (AD) is shown versus the total discord (i.e., normalized Robinson-Foulds (RF) distance between the true species tree and *estimated* gene trees), which encompasses differences due to both ILS and and gene tree estimation error (GTEE). AD is the average normalized RF distance between the true species tree and true gene trees, averaged across all genes; thus AD measures discord due solely to ILS. When GTEE is low, AD and total discord are nearly equal (points lie close to the *x* = *y* diagonal line). However, when GTEE is high, then the level of ILS is over-estimated (far below *x* = *y* line), as the total discord is much greater than the discord solely due to ILS. Each dot represents a single replicate dataset from our study (black) or representative of studies performed by Hovmöller et al. 2013 (blue) or Xi et al. 2016 (orange). Our study had much a greater range of GTEE than previous studies, and thus presented more challenging conditions for gene tree summary methods.

Characterization of the gene tree heterogeneity in four biological datasets further suggests that ILS levels can be overestimated, especially in exon datasets (Table 1). For each of these datasets, we used the mean bootstrap support of the greedy consensus tree for each gene tree as an approximate measure of the average gene tree accuracy. For all studies, the mean bootstrap support across loci is quite low – between 26% to 47% (Table 1). Under the assumption that the mean bootstrap support of these consensus trees is a reasonable approximation to the accuracy of the estimated gene trees, this suggests that gene tree accuracy is fairly low (and correspondingly that the normalized RF distance between these estimated gene trees and the true gene trees may be fairly high). This suggests that much of the discord between estimated species trees and the estimated gene trees may be due to GTEE. Similarly, the false positive rate of the bootstrap strict majority consensus trees (where all branches in the strict majority consensus tree have bootstrap support that is strictly greater than 50%) is between 11% and 45%, but the false positive rate for the greedy consensus tree is between 62% to 79% (Table 1), which also supports the hypothesis that much of the discord between estimated species tree and estimated gene trees may be due to GTEE.

Filtering genes based on mean bootstrap support increases the support of the remaining genes by 12% to 29%. Interestingly, the highly supported genes are also more similar to the the species tree computed using RAxML/ExaML on the concatenated gene alignments (Table S11). Filtering genes based on the percentage of missing data per gene greatly reduces the number of genes available, but does not result in the retained genes having greater boot-strap branch support or being more similar to the RAxML/ExaML species tree (Table S12). In fact, gene filtering based on missing data can even *decrease* the mean bootstrap support of the retained genes (e.g., see the results shown for Streicher et al. (2016) in Table S12).

## DISCUSSION

Since increases in missing data or gene tree estimation error can increase species tree error, a natural assumption has been that removing the genes with missing data or high gene tree estimation error would result in improved accuracy. Many prior studies (Jiang et al. 2014; Hosner et al. 2016; Streicher et al. 2016) have evaluated these issues on empirical (i.e., biological) datasets, and found that excluding genes in biological datasets based on missing data either had little effect or tended to decrease species tree accuracy. This decrease in accuracy prompted Jiang et al. (2014) to conclude “we see no basis for treating the exclusion of genes with missing data as a necessarily safer or more conservative approach. We see no argument for why an approach that seems to decrease accuracy under realistic conditions should be considered safer or more conservative.” The simulation study by Huang and Knowles (2016) examined the impact of filtering genes based on missing data using simulated datasets, and also found that filtering reduced accuracy – often very substantially.

On the other hand, empirical studies that deleted genes likely to have high gene tree estimation error have generally found filtering to be beneficial (Blom et al. 2017; Hosner et al. 2016; Simmons et al. 2016), although one study found it to be neutral (Chen et al. 2015). Thus, filtering genes based on missing data has generally been found to be detrimental, while filtering genes based on gene tree estimation error has sometimes been beneficial.

Our simulation study examined these issues using simulated datasets, and evaluated a larger set of methods, including four methods designed to address gene tree heterogeneity due to ILS (three of the current popular summary methods and SVDquartets) and concatenation using maximum likelihood. We also find that the impact of filtering on the basis of missing data is generally neutral to detrimental, thus confirming earlier studies. In addition, we find that the impact of filtering based on missing data depended on the model condition, with higher impact for high ILS and GTEE conditions.

Our study finds that filtering based on gene tree estimation error can be beneficial in some conditions and detrimental in others. Most importantly, we find that filtering based on gene tree estimation error is only beneficial under the low ILS condition. This observation is also consistent with prior studies, where filtering tended to be considered beneficial, and where the level of ILS was suspected to be low (Hosner et al. 2016).

More generally, we find that filtering based on either criterion can have a negative impact on species tree accuracy when the number of genes becomes too small, with larger decreases in accuracy under conditions with high GTEE and high ILS.

Our results shed light on the difference in findings from earlier studies on the impact of filtering. That is, filtering reduces the amount of data available to the species tree method, and so generally speaking should reduce accuracy. However, if the data quality improves through filtering, then there can be benefit to species tree estimation. In other words, filtering is fundamentally a question of data quality versus data quantity.

Therefore, if filtering does not improve the gene quality, there is no benefit to be had from filtering – the quantity goes down but the quality does not go up. Filtering based on missing data in our experiment led to either similar or slightly better GTEE (Fig. S12), so that the data quality either stayed the same or slightly improved; however filtering substantially reduced data quantity, making the net effect generally negative rather than beneficial. This explains why filtering based on missing data is not advisable in our simulation – the improvement in data quality (measured by average GTEE in the retained genes) was too small to offset the impact of reduction in data quantity produced by filtering.

When we filtered based on gene tree estimation error, filtering led to substantial improvements in GTEE (Fig. S8), which were much greater than the improvements observed when filtering based on missing data (Fig. S12). Thus, filtering based on gene tree estimation error had a larger impact on data quality than filtering based on missing data. Even so, the impact on species tree estimation of filtering based on gene tree estimation error was only beneficial under the low to moderate ILS condition.The explanation in this difference in performance is that for low enough ILS conditions, even a few highly accurate gene trees are sufficient to estimate the true species tree (e.g., imagine the no-ILS condition, where one perfect gene tree is identical to the species tree), and so deleting genes based on GTEE will benefit summary methods. On the other hand, when ILS is high enough, the only chance of estimating the species tree well is to have a large sample of gene trees; hence, unless the sample of genes is large enough to allow many genes to be retained after filtering, deleting genes will be detrimental. Therefore, the data quantity requirement depends not only on the method but also on the model condition, and most species tree methods probably require more data under high ILS model conditions for highly accurate species trees to be estimated (Shekhar et al. 2017).

We can now examine the prior studies with this framework in mind. Most other studies (Jiang et al. 2014; Hosner et al. 2016; Streicher et al. 2016) have concluded that filtering based on missing data is at best neutral, but can also be detrimental, e.g., decreasing the bootstrap support or recovery of well-established clades. It is worth noting that the simulation study by Huang and Knowles (2016) observed a substantial negative impact on species tree estimation. To understand this, recall that Huang and Knowles (2016) designed their study so that the deleted genes had higher rates of evolution on average than the retained genes, and were hence more phylogenetically informative than the retained genes. This suggests that the deleted genes would probably have more accurate gene trees, and that filtering based on missing data would increase the average gene tree estimation error in the retained genes compared to the original set. Hence, a likely explanation for why Huang and Knowles (2016) found that filtering based on missing data very substantially reduced accuracy is that the filtering they did *reduced* the average gene tree accuracy –— the opposite of the purpose of filtering genes!

Another example where a prior empirical study found filtering based on a proxy for gene tree error to be beneficial is Hosner et al. (2016), who found that filtering the genes to retain only the top 5% of loci with the highest number of parsimony informative sites greatly increased the topological similarity between trees estimated using summary methods (ASTRAL, ASTRID) and trees estimated using site-based methods (unpartioned RAxML and SVDquartets). Not surprisingly, filtering increased the mean branch support of the bootstrap greedy consensus trees from 28% to 57% – and the total number of genes available to summary methods was *~*240. In fact, Hosner et al. (2016) hypothesize that “ILS is relatively low… and that ML estimate of galliform phylogeny was not affected by the anomaly zone.” Thus, filtering based on this proxy for gene tree estimation error seemed to improve gene tree quality and species tree estimation, and under conditions where ILS was considered to be low –— exactly the condition in which we predict filtering should help.

## CONCLUSIONS

Our study examines four coalescent species tree estimation methods (ASTRAL, ASTRID, MP-EST, and SVDquartets) and unpartitioned concatenation using maximum likelihood (RAxML) under a wide range of model conditions. Overall, the most accurate methods are ASTRAL, ASTRID, unpartitioned concatenation using maximum likelihood (RAxML) – with the relative performance depending on the model condition. RAxML generally outperforms other methods when the level of ILS is sufficiently low and can even be more accurate than the coalescent methods under higher levels of ILS when gene tree estimation error is very high. MP-EST is generally less accurate than ASTRAL and ASTRID.

Although SVDquartets is typically among the least accurate methods, it can be more accurate than the summary methods under conditions of extremely high gene tree estimation error. Although missing data and low signal in the different gene alignments (and hence gene tree estimation error) impact all these methods (and some more so than others), the relative performance remains unchanged across different model conditions.

Gene filtering based on gene tree estimation error is observed to improve summary methods only when the level of ILS is sufficiently low and the number of genes is sufficiently high; a finding consistent with prior studies finding that gene filtering based on approximations of gene tree estimation error results in species trees more similar to those estimated by maximum likelihood methods (e.g., RAxML) on the concatenated gene alignments. Otherwise, excluding genes based on gene tree estimation error or missing data from species tree inference is likely to reduce species tree accuracy rather than improve it.

As both gene tree estimation error and incomplete lineage sorting affect the relative performance of methods and the impact of gene filtering, we recommend that systematic studies attempt to quantify these features in their dataset prior to estimating species trees. Future research is needed to develop tools to accurately quantify levels of gene tree estimation error and incomplete lineage sorting in empirical datasets.

On the positive side, the fact that filtering genes is generally not advisable (not even to filter out genes with high gene tree estimation error) shows that many current species tree estimation methods are able to extract meaningful (and useful) phylogenetic signal under very challenging conditions. In particular, summary methods - although clearly impacted by gene tree estimation error - tend to improve with additional genes, even when they have high estimation error - when attempting to construct species trees that have high ILS levels. This is encouraging, and suggests the possibility that given enough data (i.e., enough genes) very accurate species trees might be recovered using some of these methods. On the other hand, the total amount of data that might be required to reach high accuracy may be beyond what biological datasets can provide.

The generally excellent performance of concatenation, even in the simple unpartitioned approach used here, is also encouraging. But it seems unlikely that concatenation, even with partitioning, will be able to recover species trees with high accuracy under very high levels of ILS; theory argues against this and simulation studies do not support this.

So the challenge remains: can very challenging phylogenetic trees be reconstructed, in the presence of substantial ILS? Highly accurate gene trees are unlikely, since short species tree branches will produce difficult gene trees, and long recombination-free regions are unlikely to exist in deep phylogenies. Site-based methods might be the right direction, but this study does not suggest that the currently available site-based methods are as accurate as the alternative methods. Co-estimation of gene trees and species trees might be the most powerful approach, but current co-estimation methods are computationally intractable on large datasets.

In fact, one of the challenges of species tree estimation is that much of the tree might be easy to estimate but some key nodes might resist analysis. In particular, incomplete lineage sorting and gene tree estimation error are likely to both result when species trees have very short branches, producing conditions where all current approaches can fail to produce high accuracy. It is certainly clear that summary methods, which depend on gene trees, have difficulty in these cases, and concatenation is also known to fail in these cases because it does not take gene tree heterogeneity into account. It is also possible that site-based methods, such as SVDquartets, would also have difficulty resolving very short branches.

These observations suggest that the practice of mainly focusing on highly conserved regions of the genome may not be sufficient for resolution of these challenging nodes. Indeed, including regions that are faster evolving might improve species tree estimation. Certainly, phylogenetic analyses of faster evolving sequences will imply other challenges, but they may help in the recovery of the most challenging species trees. Furthermore, this study shows that species tree estimation methods benefit from the inclusion of additional genes, even those that are not present in all (or perhaps even most) of the species.

Finally, this study suggests also the need to develop new methods for species tree estimation, and to evaluate these methods under a range of conditions, including ones where gene tree estimation error is high. While it remains to be seen whether the levels of ILS that are commonly encountered in biological studies are high enough to make traditional approaches like concatenation inapplicable, certainly new methods are needed for the more challenging datasets that may be currently intractable using existing methods.

## ACKNOWLEDGEMENTS

EKM is supported by the National Science Foundation Research Fellowship Program under Grant Number DGE-1144245. TW is supported by National Science Foundation Grant Number CCF-1535977. This research is part of the Blue Waters sustained-petascale computing project, which is supported by the National Science Foundation (awards OCI-0725070 and ACI-1238993) and the state of Illinois. Blue Waters is a joint effort of the University of Illinois at Urbana-Champaign and its National Center for Supercomputing Applications. This research also made use of the Illinois Campus Cluster, a computing resource that is operated by the Illinois Campus Cluster Program in conjunction with the National Center for Supercomputing Applications and which is supported by funds from the University of Illinois at Urbana-Champaign.

